# *Leishmania major centrin* knock-out parasites alter the kynurenine- aryl hydrocarbon receptor signaling to produce a pro-inflammatory response

**DOI:** 10.1101/2022.09.15.508117

**Authors:** Timur Oljuskin, Nazli Azodi, Greta Volpedo, Parna Bhattacharya, Nevien Ismail, Shinjiro Hamano, Greg Matlashewski, Abhay R. Satoskar, Sreenivas Gannavaram, Hira L. Nakhasi

**Affiliations:** Animal Parasitic Diseases Laboratory, Agricultural Research Service, USDA, Beltsville, 20705 MD, USA; Division of Emerging and Transfusion Transmitted Diseases, CBER, FDA, Silver Spring, MD, 20993 USA; Department of Microbiology, The Ohio State University, Columbus, OH 43210, USA; Department of Parasitology, Institute of Tropical Medicine (NEKKEN), The Joint Usage/Research Center on Tropical Disease, Nagasaki University, Nagasaki, Japan and Nagasaki University Graduate School of Biomedical Sciences Doctoral Leadership Program, Nagasaki, Japan; Department of Microbiology and Immunology, McGill University, Montreal, Canada; Department of Pathology, Wexner Medical Center, The Ohio State University, Columbus, OH 43210, USA

## Abstract

Leishmaniasis is a parasitic disease that is prevalent in approximately 88 countries, and yet no licensed human vaccine exists against it. Towards control of leishmaniasis, we have developed *Leishmania major centrin* gene deletion mutant strains (*LmCen*^*-/-*^) as a live attenuated vaccine, which induces a strong Th1 response to provide IFN-γ-mediated protection to the host. However, the immune mechanisms of such protection remain to be understood. Metabolomic reprogramming of the host cells following *Leishmania*-infection has been shown to play a critical role in pathogenicity and shaping the immune response following infection. Here, we applied untargeted mass spectrometric analysis to study the metabolic changes induced by infection with *LmCen*^*-/-*^ and compared those with virulent *L. major* parasite infection to identify the immune mechanism of protection. Our data shows that immunization with *LmCen*^-/-^ parasites, in contrast to virulent *L. major* infection, alters tryptophan metabolism to down-regulate kynurenine-AhR signaling and promote a pro-inflammatory response.

## Introduction

Leishmaniasis is a parasitic disease caused by the various strains of the blood-borne protozoan species *Leishmania*, which is transmitted to humans via the bite of an infected sandfly. The disease pathology ranges from localized skin ulcers (cutaneous leishmaniasis, CL) that can deteriorate into disfiguring scars (mucocutaneous leishmaniasis, MCL) to fatal systemic disease (visceral leishmaniasis, VL), depending on the species of the infecting *Leishmania* parasite. CL is the most prevalent form of the disease, which is commonly caused by *Leishmania major* in the old world and *Leishmania mexicana* in the Americas (Kaye and Scott, 2011; Reithinger et al., 2007; Torres-Guerrero et al., 2017). Approximately, one billion people are at risk of infection, mainly in tropical and subtropical countries, and autochthonous infections have been reported in the southern regions of the United States (Curtin and Aronson, 2021); however, no licensed vaccines currently exist, and treatment options remain limited (Kaye and Scott, 2011; Torres-Guerrero et al., 2017).

Towards the control of leishmaniasis, our group has previously developed a genetically modified *L. major* mutant strain where the *centrin* gene was deleted (*LmCen*^*-/-*^) via CRISPR/cas-9 technology (Zhang et al., 2020b). These studies were based on previous investigations that showed deletion of centrin from *L. donovani* results in a significant growth defect in the amastigote form of the parasite and loss of virulence (Selvapandiyan et al., 2004). Immunization with centrin deleted *L. donovani* protected hosts against virulent challenge in studies with mice and dogs (Fiuza et al., 2015; Selvapandiyan et al., 2009). The deletion of the *centrin* gene from *L. major* similarly caused a significant growth defect in the amastigote from of the parasite and loss of virulence that resulted in a lack of skin lesions and gradual clearance of *LmCen*^*-/-*^ parasites from the host (Zhang et al., 2020b). Additionally, immunization with the *LmCen*^*-/-*^parasites induced a strong Th1response in the hosts and protected them against challenge by the virulent *L. major* or *L. donovani* strains (Karmakar et al., 2021; Zhang et al., 2020b). The *LmCen*^*-/-*^ strain has shown excellent safety and efficacy in pre-clinical studies and is currently under GMP production towards future clinical trials as a live attenuated vaccine candidate against leishmaniasis.

Several experimental *Leishmania* vaccines have been tested in pre-clinical models and a few have advanced to clinical trials (Volpedo et al., 2021). However, the immune mechanisms of protection beyond those involved in the induction of Th1 immunity remain to be explored. It has been shown that *L. major* parasites can interfere with the host’s immune signaling pathways and modulate the expression of cytokines to create an environment that benefits their growth and immune evasion (Gupta et al., 2013; Makala et al., 2011; Moreira et al., 2015). For example, IFN-γ production is critical for the destruction of intracellular parasites through the induction of nitric oxide, superoxide anions, and GTPases (Liew et al., 1990; Murray, 1981). Previous studies have established that the persistence of *Leishmania* parasites in the host are strongly dependent on the inhibition of IFN-γ production, the induction of the anti-inflammatory IL-10 and an elevated level of TGF-β that inhibit anti-microbial macrophage functions (Barral-Netto et al., 1992; Chandra and Naik, 2008; Guizani-Tabbane et al., 2004; Kaye and Scott, 2011; Nandan and Reiner, 1995). On the other hand, previous work on the efficacy of the *LmCen*^-/-^ parasites, has established that immunization with the *LmCen*^*-/-*^ strain significantly increases the pro-inflammatory Th1 response following challenge with either virulent *L. major* or *L. donovani*, indicated by the presence of IFN-γ producing effector T-cells analogous to leishmanization (Zhang et al., 2020b) (Karmakar et al., 2021). In contrast, another *centrin* deletion mutant in *L. mexicana* background also conferred protection against homologous challenge in the immunized hosts by downregulating IL-10 response, while the IFN-γ mediated Th1 response remained unchanged between the wild type and mutant parasite infections (Volpedo et al., 2020b). The immune mechanisms of protection thus appear to be different in *LmCen*^-/-^ and *LmexCen*^*-/-*^ strains.

While pathogenicity and vaccine efficacy characteristics are typically addressed from an immunological perspective in parasitic organisms, numerous studies have highlighted the importance of metabolomic reprogramming in disease development (Hu et al., 2022; O’Neill et al., 2016; Pavlova and Thompson, 2016; Ward and Thompson, 2012). In *Leishmania* infection, early differentiation of macrophages into M1 and M2 states, and the associated immune response that either favors the host or the parasite respectively, is driven by metabolic changes (Moreira et al., 2015; Naderer and McConville, 2008; Ty et al., 2019). Such metabolic reprogramming occurs prior to the onset of an immune response, highlighting the role of metabolic products as ligands for transcription factors that turn on the expression of cytokines and other immune molecules (Ganeshan and Chawla, 2014). Increased levels of glutaminolysis and oxidative phosphorylation have been shown to determine the activation status of plasmacytoid DCs (Basit et al., 2018; Thwe et al., 2017). Increased fatty acid oxidation or glycolysis causes TLR4 mediated DC activation, demonstrating the role of metabolic changes in shaping innate immune responses (Basit and de Vries, 2019; Basit et al., 2022).

Distinct metabolic reprogramming of phagocytic cells has also been demonstrated in *Leishmania* infected macrophages. It has been reported that both gluconeogenesis and the pentose phosphate pathway are necessary for amastigote replication in the macrophages (Ilg, 2002; Maugeri et al., 2003; Naderer et al., 2006; Naderer and McConville, 2008), where fatty acids (Hart and Coombs, 1982; Naderer and McConville, 2008; Rosenzweig et al., 2008) and amino acids (Geraldo et al., 2005; Naderer and McConville, 2008; Shaked-Mishan et al., 2006) can serve as vital carbon sources for the parasites. Similarly, *L. infantum* parasites manipulate the AMPK pathway to significantly increase the oxidative phosphorylation necessary for amastigote replication (Moreira et al., 2015).

While these studies have established the role of metabolic programming in *Leishmania* pathogenesis, similar studies addressing the vaccine-mediated immune response in *Leishmania* have not been performed. Studies in viral vaccines such as ΔF/TriAdj, a vaccine against Respiratory Syncytial Virus (RSV) showed that immunization with the attenuated viruses affects tryptophan metabolism, while virulent RSV infection primarily impacts amino acid biosynthesis and the Urea cycle (Sarkar et al., 2019).

We undertook studies to explore the metabolic pathways underlying the immune mechanisms of protection associated with *LmCen*^*-/-*^ immunization in mice. An untargeted metabolomics analysis via LC/MS was performed followed by pathway enrichment and integrative network analysis. Our results showed various metabolic pathways were significantly enriched in *LmCen*^*-/-*^ immunization, which could be driving the immune-protective benefits of the vaccine. In an accompanying study (Volpedo et al., 2022), we performed similar metabolomics analysis of mice immunized with *LmexCen*^*-/-*^ strain, which showed that the underlying metabolic drivers of immune protection are distinct in the two vaccine candidates.

## RESULTS

### Mass spectrometric analysis of the ear tissue

The ear tissues of mice intradermally inoculated with either *LmWT, LmCen*^-/-^, or PBS as a control were used to perform mass spectrometry to identify metabolites and metabolomic pathways associated with the immune mechanisms that underlie infection and immunization (Figure 1A). A two-way comparison between *LmWT* vs Naïve, *LmCen*^-/-^ vs Naïve, and *LmCen*^-/-^vs *LmWT* were performed to identify metabolites associated with the immune mechanism of pathogenesis or protection. The two-way comparison files contained 4610, 5158, and 3676 features, respectively. Volcano plots with a fold change threshold of ±2 and a *p-* value threshold of 0.05 were used to identify the significant features in the datasets for both the positive (Figure 1B) and the negative (Figure 1C) modes of the mass spectrometry with the former identifying the protonated and the latter identifying the non-protonated molecules. To visualize the overall structure of the datasets and determine whether unique metabolite signatures exist among the three comparisons, Partial-Least Squares Discriminant (PLSD-DA) analysis, a multivariate dimensionality reduction method, was performed using the data from both positive (Figure 1D) and the negative (Figure 1E) modes. The PLSD-DA clustered the metabolites into three non-overlapping groups with 17.7%-34.7% and 4.6%-8.6% of the variance explained by the first two principal components, respectively, indicating that the three conditions of naïve, virulent infection and immunization lead to distinct enrichments.

**Figure 1.**
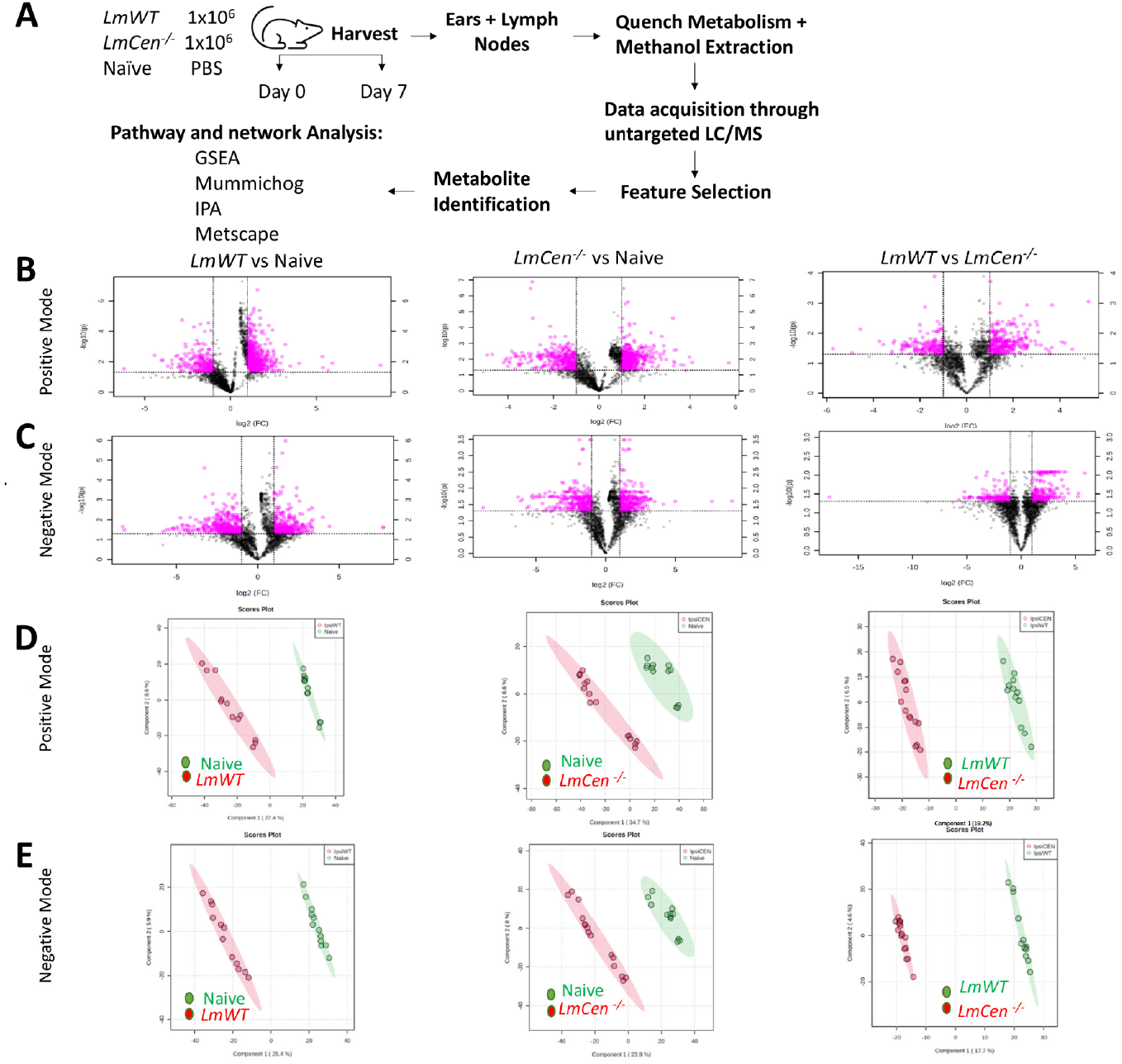
Infection with *LmWT* and immunization with *LmCen*^*-/-*^ display distinct metabolic signatures at the inoculation site. Female C57Bl/6 mice (n=≥4 per group) were injected intradermally in the ear with stationary phase *LmWT, LmCen*^*-/-*^ parasites. After 7 days, the ear tissues were collected and processed for mass spectrometry. The data acquired was used to identify enriched metabolites and construct activated metabolomic pathways **(A)**. Normalized data from ear tissue of C57Bl/6 mice was used to perform statistical analysis **(B, C)**. Features selected by volcano plot with < 0.05 false-discovery rate (FDR) and > 2-fold change (FC) threshold cutoffs from positive **(B)** and negative **(C)** modes for ear tissue of *LmWT* vs. naïve, *LmCen*^*-/-*^ vs. naïve, and *LmCen*^*-/-*^ vs. *LmWT* mice with log-transformed fold change (x) 2 and t-tests thresholds (y) 0.05 **(D, E)**. Partial least squares-discriminant analysis (PLS-DA) from positive **(D)** and negative **(E)** mode for ear tissue of *LmWT* vs. naïve, *LmCen*^*-/-*^ vs. naïve, and *LmCen*^*-/-*^ vs. *LmWT* mice indicating the existence of distinct metabolic signatures under the infection and immunized conditions. Abbreviations: LC/MS: liquid chromatography / mass spectrometry; GSEA: gene set enrichment analysis.

### Arginine metabolism is significantly altered in *LmCen*^*-/-*^ immunization compared to *LmWT* infection

To identify metabolic pathways distinctly enriched in the ear tissues immunized with *LmCen*^*-/-*^ in comparison to the naïve and the *LmWT*-infected mice tissues, both the mummichog and the Gene Set Enrichment Analysis (GSEA) algorithms were used to analyze the positive (Figure 2A, 2B, and2C) and negative mode datasets (Figure 2D, 2E, and 2F). For this analysis *p*-values were integrated together via a Fisher’s method and visualized the results with MS Peaks to Pathways plots. The metabolic pathways enriched in the three different groups were identified through the comparisons of *LmWT* vs Naïve (Figure 2A and2D), *LmCen*^*-/-*^ vs Naïve (Figure 2B and 2E), and the *LmWT* infection against the *LmCen*^*-/-*^ immunization (Figure 2C and2F), indicating the distinct pathways enriched under each condition. Results showed that arginine biosynthesis to be one of the most highly enriched pathways in the *LmCen*^*-/-*^ immunization in comparison to the naïve and the *LmWT* infection (highlighted by the black box, Figure 2B). To verify the results, pathway enrichment was also conducted by Ingenuity Pathway Analysis (IPA) to identify the canonical pathways (Supplementary Figure 1). The results from IPA also showed that arginine biosynthesis is one of the most highly enriched pathways in the *LmCen*^*-/-*^ immunization in comparison to the naïve and *LmWT* datasets. Metabolites involved in the arginine biosynthesis pathway detected in the normalized LC/MS outputs and their fold change levels are listed (Table 1). Our data showed that L-arginine is enriched in both the positive mode *LmWT* infection and the *LmCen* ^*-/-*^ immunization in comparison to the naïve control. However, the *LmWT* vs *LmCen*^*-/-*^ comparison showed that L-arginine is more enriched in the *LmCen*^*-/-*^ compared to *LmWT* infection (Table 1). Spermine, a downstream product of the arginine metabolism derived from spermidine, appeared only in the *LmCen*^*-/-*^ dataset but not in *LmWT*. Spermine was highly enriched in both the *LmCen*^*-/-*^ vs naïve and *LmCen*^*-/-*^ vs *LmWT* comparisons (Table 1).

**Figure 2.**
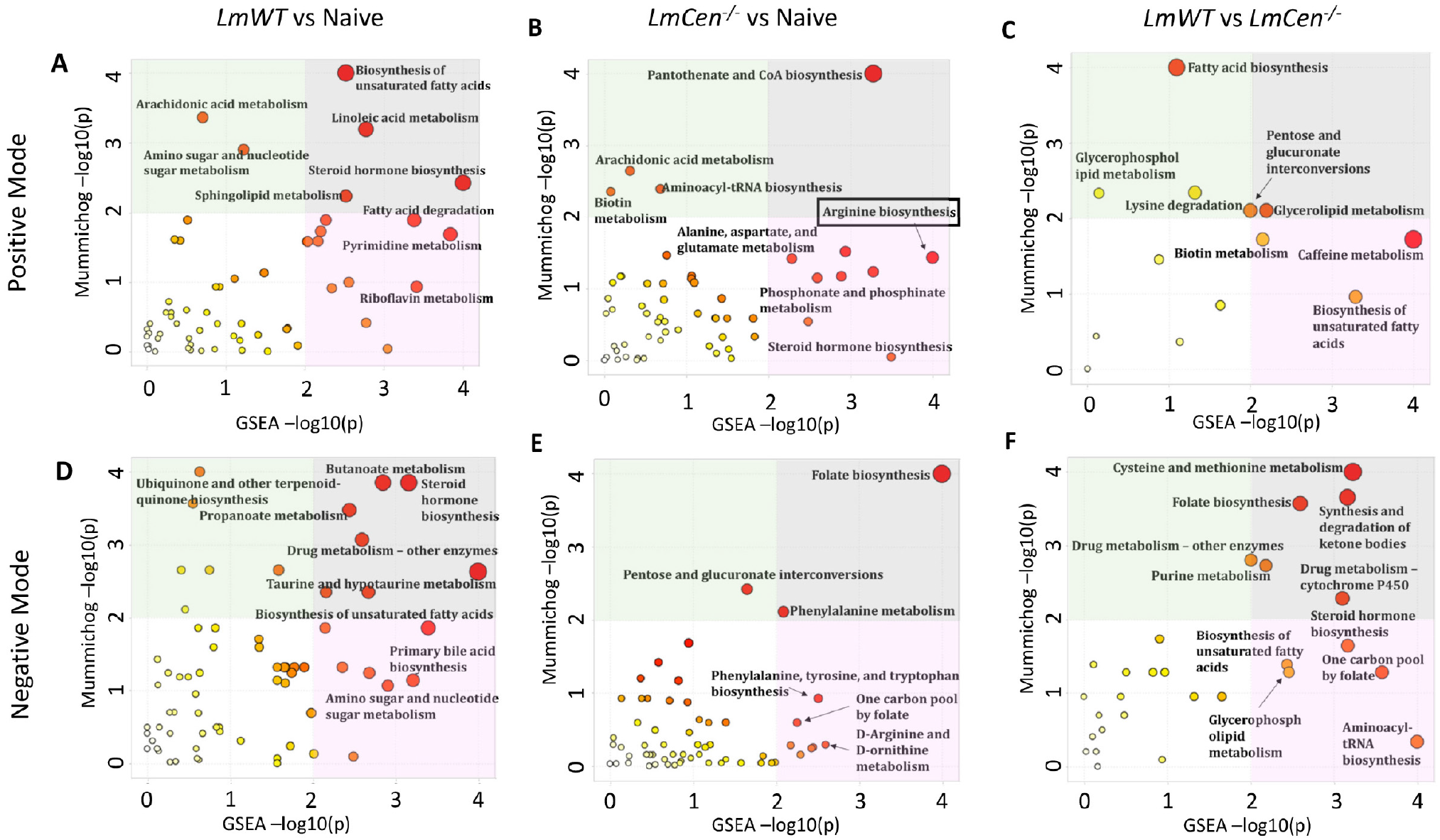
Metabolic pathways enriched in mouse ear tissues infected with *LmWT* or immunized with *LmCen*^*-/-*^. Normalized data from ear tissue of C57Bl/6 mice after 7 days of infection with *LmWT*, immunization with *LmCen*^*-/-*^ or naïve control was used to perform peaks to pathway analysis. Using the MS Peaks to Paths module in MetaboAnalyst5.0, the mummichog and gene set enrichment analysis (GSEA) p-values were combined. The Integrated MS Peaks to Paths plot summarizes the results of the Fisher’s method for combining mummichog (y) and GSEA (x) p-values from the positive **(A)** negative **(B)** mode data sets, indicating the metabolic pathways enriched. The size and color of the circles correspond to their transformed combined p-values. Large and red circles are considered the most perturbed pathways. The colored areas show the significant pathways based on either mummichog (green) or GSEA (pink), and the gray area highlights significant pathways identified by both algorithms. Arginine metabolism pathway associated with a protective immune response was found to be enriched in *LmCen*^*-/-*^immunization (highlighted by the black boxes).

**Table 1.** List of significant metabolites of the arginine metabolism pathway detected by LC/MS and their respective enrichment levels in mice infected with *LmWT* or immunized with *LmCen*^*-/-*^. Normalized data from ear tissue of C57Bl/6 mice after 7 days of infection with *LmWT*, immunization with *LmCen*^*-/-*^ or naïve control was used to analyze the concentration levels of metabolites associated with arginine metabolism that were detected in our dataset by mass spectrometry. Different isomers of metabolites could be detected multiples times through various m/z ratios recognized by the mass spectrometry analysis. Metabolites shown in italics were detected by the negative mode LC/MS, while the remainder were detected by the positive mode.

| Dataset | Compound | m/z Ratio | Log FC |
| --- | --- | --- | --- |
| *LmWT* vs Naïve | L-Arginine | 175.120515789341/6.53941666666667 | 0.711983 |
|  |  | 174.111888273963/12.3793333333333 | 0.175912 |
|  | L-Ornithine | 155.079258810298/4.28323333333333 | 0.777981 |
|  | L-Citrulline | 198.085631129694/1.26938333333333 | 0.143353 |
|  |  | 176.102936662285/1.26938333333333 | 0.144771 |
|  |  | 176.1017753953/8.26498333333333 | Infinity |
|  |  | 351.196228027344/6.74226666666667 | -1.53448 |
|  | *L-Arginine* | 347.218973727799/17.9363333333333 | -0.85208 |
|  | *L-Citrulline* | 174.087391981391/1.2694 | 0.129864 |
|  |  | 175.094959488147/8.52898333333333 | 1.871527 |
| *LmCen*<sup>-/-</sup> vs Naïve | L-Arginine | 175.117476556853/6.35535 | 0.191052 |
|  | L-Citrulline | 175.09342956543/14.2167833333333 | -0.40031 |
|  |  | 198.085631129694/1.26938333333333 | 0.075425 |
|  |  | 176.102936662285/1.26938333333333 | 0.064003 |
|  |  | 351.198661460237/5.61805 | -0.17327 |
|  | Spermine | 241.17788404089/12.7144666666667 | 0.032859 |
| *LmWT* vs *LmCen*<sup>-/-</sup> | L-Arginine | 174.111485044507/10.6618166666667 | 0.413899 |
|  |  | 175.117476556853/6.35535 | -0.19811 |
|  | L-Citrulline | 176.102936662285/1.26938333333333 | 0.080768 |
|  |  | 198.085631129694/1.26938333333333 | 0.067928 |
|  | Spermine | 241.178182435098/11.1409833333333 | -0.20232 |
|  | *L-Arginine* | 209.080528261553/2.07785 | -0.40848 |
|  |  | 347.218973727799/17.9363333333333 | -0.99784 |
|  | *L-Citrulline* | 174.087391981391/1.2694 | 0.067751 |
|  |  | 175.094959488147/8.52898333333333 | 1.173381 |
|  |  | 210.065612776268/7.64563333333333 | Infinity |

### Polyamines that enable parasite proliferation are enriched in the *LmWT* infection

Arginine and citrulline are metabolites connected in a looped relationship in which citrulline is a byproduct of the nitric oxide production via arginine metabolism, and it can be used in arginine synthesis as well through the ornithine cycle. Accordingly, the results of IPA also showed that citrulline metabolism was highly enriched in the *LmWT* infection in comparison to the naïve and the *LmCen*^*-/-*^ infection (Supplementary Figure 1). Ornithine was exclusively enriched in the *LmWT* vs naïve comparison and was not detected in the *LmCen*^*-/-*^ vs naïve datasets (Table 1), indicating its importance of polyamines to the parasite proliferation.

### Kynurenine levels are altered between *LmWT* infection and *LmCen*^*-/-*^ immunization

Previous studies have shown that tryptophan metabolism is often altered in conjunction with arginine metabolism through the interaction of Arg1 and indoleamine 2,3-dioxygenase 1 (IDO1) (Crowther and Qualls, 2020; Mondanelli et al., 2017; Mondanelli et al., 2019). Tryptophan can be catabolized through the three distinct mechanisms to produce kynurenine, indole, and serotonin (Hsu et al., 2020). In the kynurenine pathway, IDO1 acts as the rate-limiting enzyme in the conversion of the substrate tryptophan into kynurenine and other downstream metabolites such as kynurenic Acid. In the indole pathway, Indole-3-acetaldehyde can yield 6-formylindolo[3,2-*b*]carbazole (FICZ), while the serotonin pathway leads to the production of melatonin (Hsu and Tain, 2020; Mondanelli et al., 2019). Kynurenine, FICZ and melatonin have been shown to have immunoregulatory roles and are of interest in the context of vaccine immunity.

Metabolites associated with the tryptophan metabolism pathway were searched for in the normalized LC/MS outputs to identify their enrichment. As shown in the Table 2, the *LmWT* infection is characterized by a depletion of tryptophan and higher levels of Indole-3-acetaldehyde and one isoform of kynurenine, while two kynurenine isoforms showed a slight downregulation. In contrast, the *LmCen*^*-/-*^ immunization is distinguished by higher levels of tryptophan, reduced level of Indole-3-acetaldehyde and a downregulation of the kynurenine. Interestingly, melatonin was found highly enriched in the *LmCen*^*-/-*^ dataset but was undetected in the *LmWT* infection (Table 2).

**Table 2.** List of significant metabolites of the tryptophan pathway detected by LC/MS and their respective enrichment levels in mice infected with *LmWT* or immunized with *LmCen*^*-/-*^. Normalized data from ear tissue of C57Bl/6 mice after 7 days of infection with *LmWT*, immunization with *LmCen*^*-/-*^ or naïve control was used to analyze the concentration levels of metabolites associated with tryptophan metabolism pathway that were detected in our dataset by mass spectrometry. Different isomers of metabolites could be detected multiples times through various m/z ratios recognized by the mass spectrometry analysis.

| Dataset | Compound | m/z Ratio | Log FC |
| --- | --- | --- | --- |
| *LmWT* vs Naïve | Tryptophan | 409.183727323844/16.2527 | -0.84285 |
|  | Indole-3-acetaldehyde | 160.074993175059/8.60266666666667 | 0.412018 |
|  | Kynurenine | 417.177374622835/6.63973333333333 | -0.5874 |
|  |  | 209.091836598665/3.50861666666667 | -0.08652 |
|  |  | 209.094131469727/3.25915 | 0.354958 |
| *LmCen*<sup>-/-</sup> vs Naïve | Tryptophan | 205.098835651367/1.6815 | 0.202931025 |
|  | Indole-3-acetaldehyde | 160.075205801606/4.96223333333333 | 0.106710051 |
|  |  | 160.075511683185/7.90605 | 0.250066463 |
|  | Kynurenine | 417.177374622835/6.63973333333333 | -0.715094839 |
|  | Melatonin | 232.120675751645/6.94911666666667 | 6.138261 |
|  | Kynurenic Acid | 190.049071881504/7.86811666666667 | Infinity |
| *LmWT* vs *LmCen*<sup>-/-</sup> | Tryptophan | 205.098835651367/1.6815 | -0.52385 |
|  |  | 204.08733700045/14.1969 | Infinity |
|  |  | 205.097443451686/4.99411666666667 | -0.07056 |
|  | Indole-3-acetaldehyde | 160.075205801606/4.96223333333333 | -0.08637 |
|  | Kynurenine | 209.091836598665/3.50861666666667 | -0.15887 |
|  |  | 417.177401765291/5.18043333333333 | -0.27253 |

To better visualize the alterations in tryptophan metabolism, compound-gene network analysis of *LmCen*^*-/-*^ vs naïve and *LmWT* vs naïve was performed and is shown in Figures 3A and 3B, respectively. This analysis takes into account the variations in the fold changes of isoforms reported in Table 2. Results showed that the tryptophan was detected in both *LmCen*^-/-^ and *LmWT* infections although reduced in the *LmWT* infection, while the products of tryptophan catabolism, i.e., kynurenine and indole-3-acetaldehyde were significantly reduced in *LmCen*^*-/-*^ immunization compared to *LmWT* infection (Figure3A and 3B). In contrast, melatonin was enriched in *LmCen*^*-/-*^ infection but was not detected in *LmWT* infection (Figure 3A and 3B).

**Figure 3.**
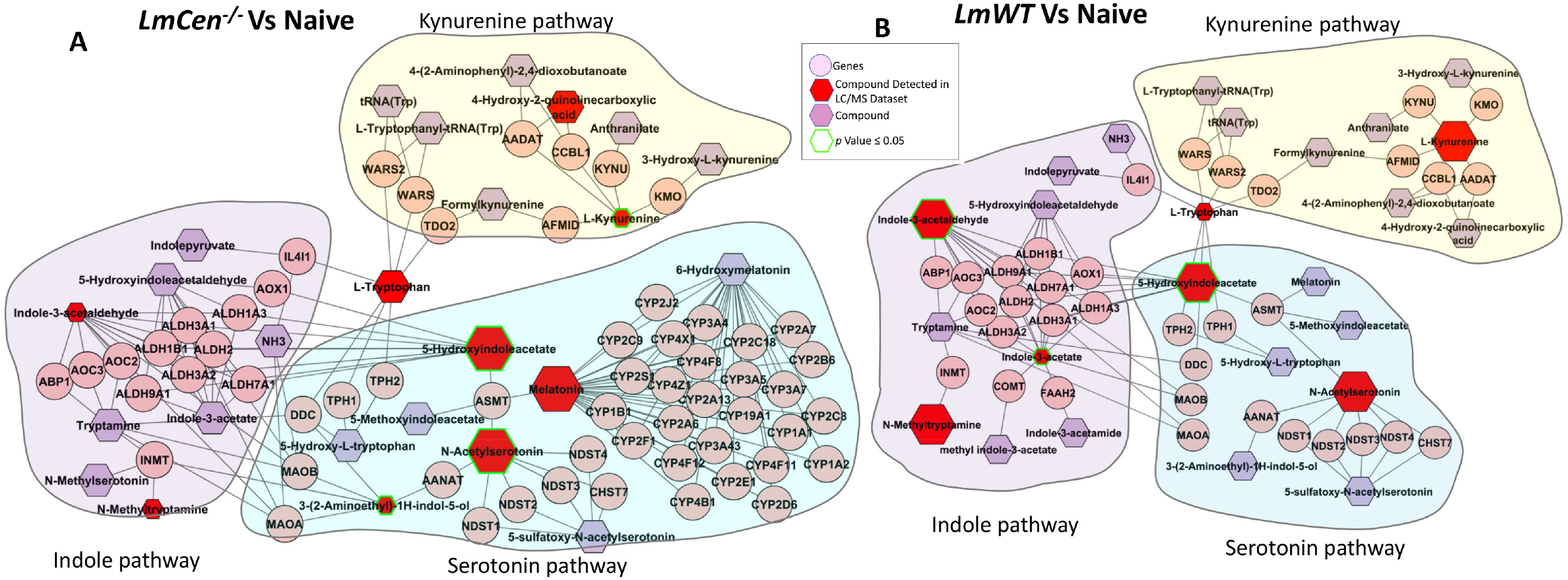
Infection with *LmWT and LmCen*^*-/-*^ lead to differential regulation of the tryptophan metabolism pathway. Normalized data from the ear tissue of C57Bl/6 mice infected with *LmWT*, immunized with *LmCen*^*-/-*^ and naïve controls was used to perform integrative compound-compound network analysis with Metscape. The differential regulation of the tryptophan metabolites, specifically kynurenine, indole-3-acetaldehyde, and melatonin are illustrated in this graph for the immunized **(A)** and infected **(B)** conditions compared to naïve control. Larger hexagons represent up-regulation, while smaller hexagons represent down-regulation. Red hexagons represent compounds detected in the data set, while hexagons with a green outline represent statistically significant metabolites (p-value ≤ 0.05). The purple hexagons represent compounds that are associated with the pathway but are not detected in the input dataset. The pink circles represent the genes regulating the biosynthetic activities. The networks associated with the kynurenine, indole, and serotonin pathways are highlighted with yellow, purple, and blue hues respectively.

### Inhibition of kynurenine-AhR signaling by 1-methyl tryptophan increases the expression of AhR and IFN-γ

Kynurenine has been well-established in playing an immunomodulatory role through its activity as a ligand for the aryl hydrocarbon receptor (AhR) (Mezrich et al., 2010). The high abundance of kynurenine found in *LmWT* infection may thus predispose the infected cells to produce TGF-β and other anti-inflammatory cytokines (Proietti et al., 2020). To test the hypothesis that the interaction between kynurenine and AhR played a significant role in the immunity of *LmCen*^*-/-*^, 1-methyl tryptophan (1-MT), a well-characterized IDO-1 inhibitor (Wirthgen et al., 2020), was used to deplete the production of kynurenine in naïve, *LmWT* and *LmCen*^*-/-*^ infected BMDCs (Figure 4A). Gene expression levels of IDO-1, AhR, and IFN-γ were measured following IDO-1 inhibition using RT-PCR (Figures 4B and 4C). The results show that among the untreated samples, the gene expression levels of IDO-1, AhR, and IFN-γ are detected at a higher level in the *LmCen*^*-/-*^ infection in comparison to the *LmWT* infection. However, upon the addition of 1-MT, both AhR and IFN-γ gene expression levels increased in the *LmWT* (Figure 4C) and to a lesser extent in the *LmCen*^*-/-*^ infection (Figure 4B) in comparison to the untreated control. This experiment was performed twice, and an average representative is depicted in Figure 4B and 4C. The raw Ct-values are presented in the supplementary information (Supplementary Table 1).

**Figure 4.**
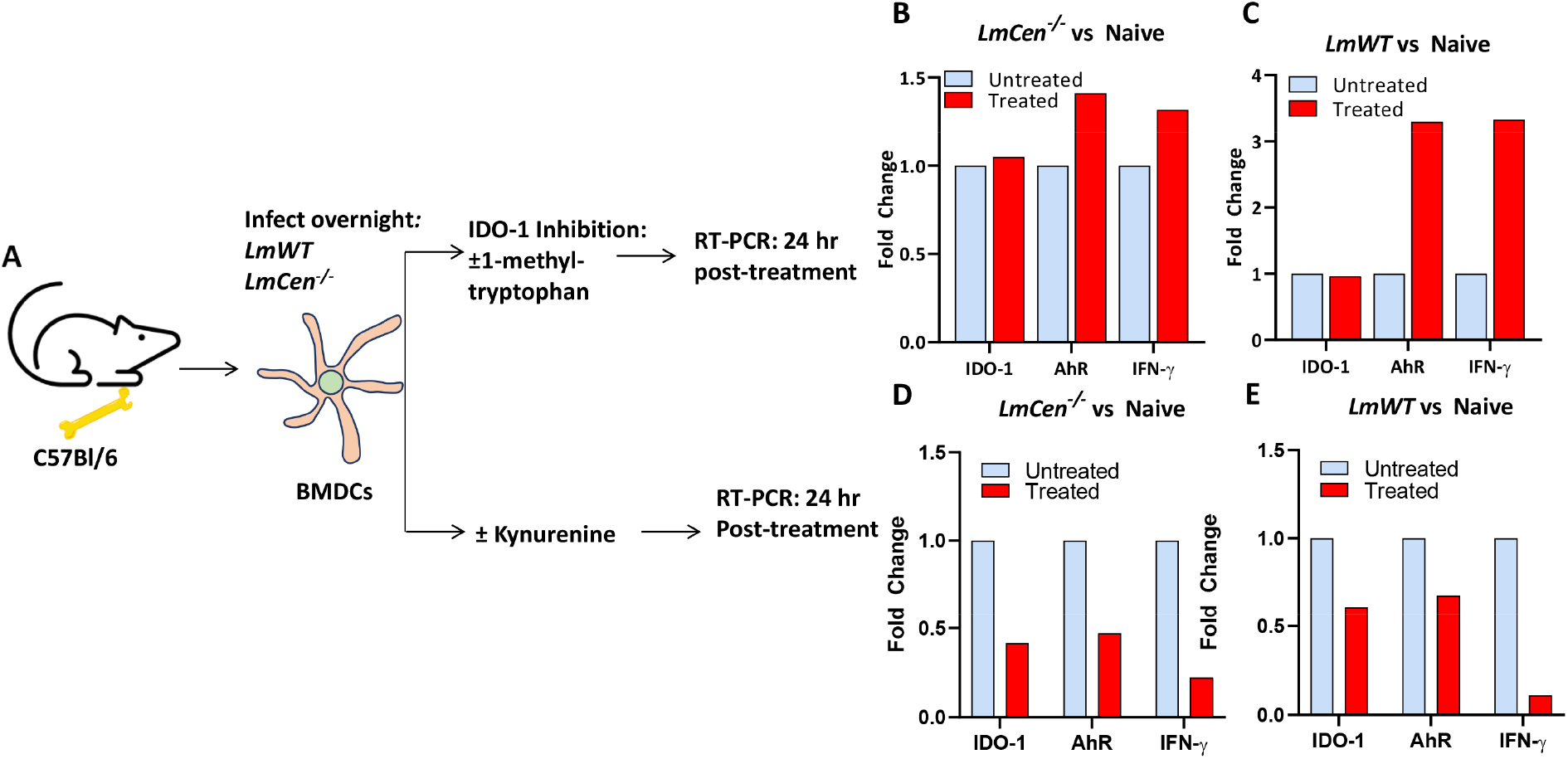
Validation of the role of tryptophan metabolism in vaccine induced immunity. To confirm kynurenine-dependent alteration of IDO-1, AhR, and IFN-γ expression in *LmWT* and *LmCen*^*-/-*^ infections, BMDCs were infected overnight with *LmWT* or *LmCen*^*-/-*^ **(A)**. After 24 hours, 1-methyl tryptophan or kynurenine were added, and the expression levels of IDO-1, AhR, and IFN-γ were measured by RT-PCR. Upon the addition of 1-MT, in both *LmCen*^*-/-*^ **(B)** and the *LmWT* **(C)** infections, the expression of AhR and IFN-γ were increased. Both *LmCen*^*-/-*^ **(D)** and *LmWT* **(E)** infections showed a decrease in AhR and IFN-γ expression upon the addition of kynurenine, showing that the expression of AhR and IFN-γ were kynurenine-dependent.

### Addition of kynurenine decreased the expression of IDO-1, AhR and IFN-γ

The AhR-kynurenine interaction induces the increased transcription of the anti-inflammatory TGF-β, and in turn reduced IFN-γ expression. To confirm that the Kyn-AhR interaction alters IFN-γ gene expression by the levels of kynurenine, we performed in-vitro experiment in which L-kynurenine was exogenously added to naïve, *LmWT* and *LmCen* ^*-/-*^ infected BMDCs (Figure 4A), after which the gene expression levels of IDO-1, and IFN-γ were measured via RT-PCR. The results showed that exogenous addition of kynurenine resulted in a reduced expression of IDO-1, and IFN-γ in the *LmCen*^*-/-*^ (Figure. 4D) and the *LmWT* infected BMDC cultures (Figure 4E). This experiment was performed twice, and an average representative is depicted in Figure 4D and 4E. Raw Ct values are presented in the supplementary information (Supplementary Table 2). A graphical summary of the results illustrating the impact of kynurenine and 1-methyl tryptophan on the gene expression levels of IDO-1, AhR and IFN-γ in the *LmWT* and the *LmCen*^*-/-*^ environments are shown (Figures 5A and 5B).

**Figure 5.**
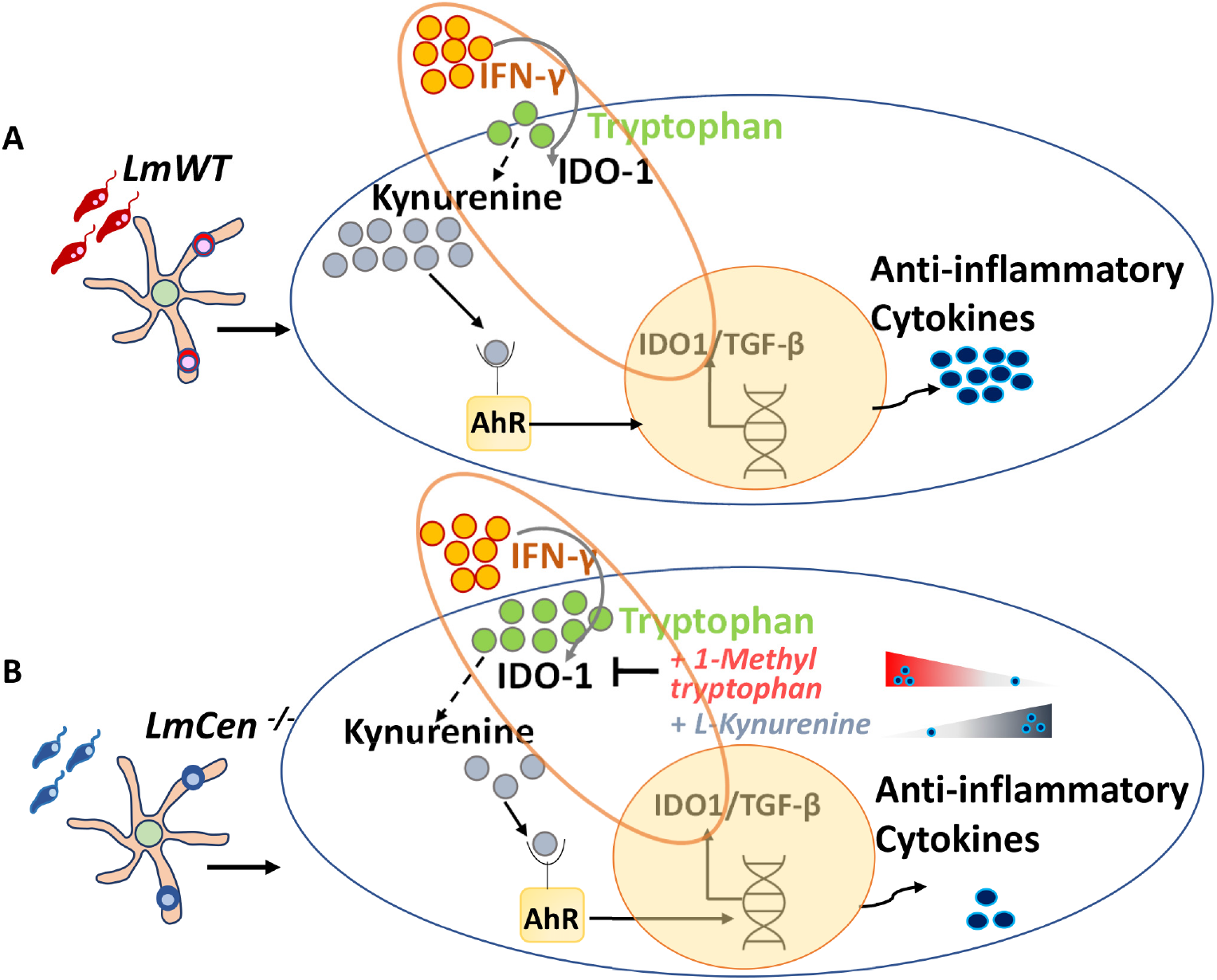
Graphical summary showcasing the kynurenine regulation and its immune-modulation in the *LmWT* infection (A) and the *LmCen*^*-/-*^ (B) immunization. *LmWT* infection causes **(A)** the tryptophan depletion, and elevated levels of kynurenine, leading to the increased anti-inflammatory cytokine response through the kynurenine-AhR signaling. In the *LmCen*^*-/-*^ immunization **(B)**, the kynurenine production is inhibited, leading to the decreased activity of AhR and anti-inflammatory signals. The use of the IDO-1 inhibitor, 1MT, led to an increased expression of IFN-γ, while the addition of kynurenine significantly decreased the IFN-γ expression.

## DISCUSSION

The live attenuated *LmCen*^*-/-*^ strain showed potent protection while maintaining the safety characteristics in pre-clinical animal models and is being developed as a candidate vaccine against leishmaniasis (Karmakar et al., 2021; Zhang et al., 2020b). Immunization with *LmCen*^-/-^ led to an enhanced Th1 response in the host in contrast to *L. major* wildtype infection (Karmakar et al., 2021; Zhang et al., 2020b). The protection following challenge in *LmCen*^-/-^ immunized hosts was mediated by the IFN-γ secreting CD4^+^ T effector cells (Zhang et al., 2020b). However, mechanisms underlying the induction of protective immunity upon immunization with *LmCen*^-/-^ parasites are not well understood (Volpedo et al., 2022a). The interaction between the host metabolism and the immune regulation that occurs in the context of an infection may provide a window to identify the mechanisms of protection and pathogenicity.

Studies of *Leishmania* infected phagocytic cells have demonstrated that metabolic changes play an important role in sustaining parasite growth through nutrient acquisition, and pathogenesis by altering of the host immune system. For example, *Leishmania* parasites are purine auxotrophs and dependent on the host’s purine metabolism to scavenge purines (Boitz et al., 2012; Marr et al., 1978). In addition to nutrient acquisition, metabolic reprogramming can also alter cytokine signaling to favor parasite growth, such as the surge in mTORC1 activity in *Leishmania*-infected cells, which leads to increased glycolysis, fatty acid biosynthesis, and oxidative phosphorylation, pathways that collectively promote parasite growth (Kumar et al., 2018; Saunders and McConville, 2020; Weichhart et al., 2015).

In addition to studies of pathogenesis, metabolomic analyses have been applied to identify immune mechanisms of protection in several vaccines. For example, metabolomic analysis to study the mechanisms of immune protection of a vaccine against *Vibrio alginolyticus* demonstrated that elevated TCA cycle activity is crucial for vaccine immunity with the intermediate metabolite, Furmate, increasing the transcription of the innate immune genes *il1β, il-8*, and *lysozyme* to promote a pro-inflammatory environment in the host (Yang et al., 2021). Similar studies in the CoronaVac (Sinovac) vaccine against Coronavirus reported that amino acid metabolism, such as phenylalanine, glycine, serine, proline, and arginine were significantly upregulated in the vaccinated hosts and played a role in the antibody production (Wang et al., 2022). Given the role that metabolic reprogramming plays in both pathogenesis and vaccine immunity, exploring such pathways may uncover the *LmCen*^*-/-*^ immunization induced mechanisms of protection.

The results of our pathway analysis showed that distinct metabolic pathways were enriched in the virulent infection verses *LmCen*^-/-^ immunization. Arginine metabolism was upregulated in both *LmWT* and *LmCen*^-/-^ infections compared to naïve controls with different products of arginine catabolism enriched in the two conditions. Arginine metabolism yields nitric oxide as one of its end-products that has anti-parasitic activity or polyamines such as ornithine and spermine that promote parasite growth (Liew et al., 1990; Naderer and McConville, 2008). Arginine metabolism can also affect the M1 and the M2 differentiation of macrophages (Gogoi et al., 2016; Naderer and McConville, 2008; Rath et al., 2014). In presence of a pro-inflammatory environment with high levels of IFN-γ, arginine metabolism leads to the synthesis of nitric oxide via nitric oxide synthase, with citrulline also being made as a byproduct that can be used for arginine synthesis as well (Gogoi et al., 2016; Rath et al., 2014). On the other hand, under an anti-inflammatory milieu, arginase can transform arginine into ornithine as a part of the urea cycle, which eventually leads to the production of proline and polyamines (Gogoi et al., 2016; Iniesta et al., 2005; Rath et al., 2014). Ornithine can be a source of nutrients for parasite; hence, elevated levels of ornithine is a characteristic of the M2 macrophages and a cellular environment that promotes the parasitic growth (Naderer and McConville, 2008). Consistent with these results, enrichment of ornithine in the *LmWT* infection suggests that arginine metabolism is being used to synthesize high levels of polyamines to support the parasite amplification (Naderer and McConville, 2008).

Interestingly, both *LmWT* and *LmCen*^*-/-*^ immunization induced an enrichment of arginine and citrulline; however, no ornithine was detected in LmCen-/- immunization.Previous reports showed that immunization with *Leishmania donovani centrin* deletion mutants induces significantly elevated nitric oxide production mediated by arginase1 enzyme than the virulent infection (Bhattacharya et al., 2015; Bhattacharya et al., 2022). It is likely that arginine metabolism produces nitric oxide in *LmCen*^-/-^ infection as well however that remains to be tested. Thus, even though arginine metabolism is enriched in both the *LmWT* and the *LmCen*^*-/-*^ infections, the products of this metabolism are distinct under the two conditions. Only spermine, a downstream product of arginine metabolism, was detected in *LmCen*^-/-^ infection. Interestingly, spermine has been shown to inhibit the induction of nitric oxide synthase (Pessenda and da Silva, 2020; Wanasen and Soong, 2008). Thus, the role of spermine in the *LmCen*^*-/-*^ immunization remains to be explored.

Arginine and tryptophan metabolisms are mechanistically linked through the activities of the rate limiting enzymes Arg1 and IDO1 (Crowther and Qualls, 2020; Mondanelli et al., 2017; Mondanelli et al., 2019). Tryptophan metabolism can also affect macrophage polarization (Naderer and McConville, 2008; Rodrigues et al., 2019). Tryptophan is typically catabolized via three metabolic pathways: the kynurenine, the indole, and the serotonin pathways. While the kynurenine (Fiore and Murray, 2021; Gargaro et al., 2016; Moffett and Namboodiri, 2003; Proietti et al., 2020; Sorgdrager et al., 2019) and indole pathways (Gargaro et al., 2016; Rannug, 2022; Sadik et al., 2020; Zhang et al., 2020c) typically induce anti-inflammatory and immunosuppressive effects, the serotonin pathway produces melatonin, known as an immune booster (Calvo et al., 2013; Carrillo-Vico et al., 2013) and a vaccine-adjuvant (Wang et al., 2021; Zhang et al., 2020a). Tryptophan metabolism was detected in both the *LmCen*^*-/-*^ and the *LmWT* datasets, however, analysis of the metabolites showed that tryptophan metabolism was differentially regulated in the immunized and infected conditions as revealed by the concentrations of kynurenine, indole-3-acetaldehyde and melatonin.

In the *LmWT* infection, the kynurenine levels were highly enriched, compared to *LmCen*^*-/-*^ immunization. Kynurenine has been the subject of much interest due to its immunomodulatory role through an interaction with the Aryl Hydrocarbon Receptor (AhR) (Mezrich et al., 2010; Nguyen et al., 2010). In this pathway, IFN-γ strongly induces the conversion of tryptophan into kynurenine, mediated by the IDO-1 enzyme (Duan et al., 2014; Fiore and Murray, 2021; Proietti et al., 2020). The kynurenine may then induce an immunomodulatory impact by binding to AhR (Clement et al., 2021; Grohmann and Puccetti, 2015; Head and Lawrence, 2009; Mezrich et al., 2010). Once activated upon the binding of kynurenine, AhR translocates from the cytoplasm into the nucleus and initiates the transcription of the anti-inflammatory TGF-β and in turn causes the reduced expression of IFN-γ (Proietti et al., 2020; Stockinger et al., 2014). This interaction creates a positive loop which results in the increased transcription of IDO-1 and an inhibition of IFN-γ, leading to higher levels of kynurenine-AhR signaling through the elevated levels of kynurenine (Munn and Mellor, 2013; Proietti et al., 2020).

The Kynurenine-AhR signaling has long been studied for its impact on the immune response to infections and cancer immunity (Stevens et al., 2009). For example, Kynurenine-AhR signaling has been investigated towards suppressing the anti-tumor immune response and promoting tumor growth (Gunther et al., 2020; Opitz et al., 2011; Prendergast, 2011; Xue et al., 2018), yielding IDO-1 inhibitors as promising candidates for cancer therapy (Diray-Arce et al., 2022; Vacchelli et al., 2014; Zhai et al., 2015). On the other hand, Kynurenine-AhR signaling has been found to generate regulatory T-cells and inhibiting the Th1 response (Campesato et al., 2020; Fallarino et al., 2006) in viral (Adu-Gyamfi et al., 2019; Fox et al., 2013; Mehraj and Routy, 2015) and parasitic (Colvin and Joice Cordy, 2020; Cordy et al., 2019; Dos Santos et al., 2020; Majumdar et al., 2019; Schmidt and Schultze, 2014) diseases, including Malaria, *Toxoplasma gondii*, and Influenza.

Previous research has shown that elevated IDO-1 expression in *L. major* infection attenuates the pro-inflammatory immune responses in a manner that leads to chronic inflammation and increased parasitic burden in the host (Makala et al., 2011). It has also been found that the kynurenine-AhR signaling in *Leishmania*-infected dendritic cells, induced elevated generation of regulatory T-cells, inhibited IFN-γ expression, and increased levels of IL-10 signaling, creating an environment that promoted parasitic growth (Nguyen et al., 2010). In agreement with these observations, our PCR data showed that the depletion of L-kynurenine by treating with 1-Methyl tryptophan, an inhibitor of IDO1 (Wirthgen et al., 2020), increased the expression of IFN-γ, while the addition of L-kynurenine significantly inhibited the IFN-γ expression presumably by acting through AhR. Taken together, these results show that the reduction of the kynurenine levels in the *LmCen*^*-/-*^ immunization may promote immunogenicity by inhibiting anti-inflammatory TGF-β and consequently promoting IFN-γ response.

Indole-3-acetaldehyde, another product of tryptophan metabolism was enriched in the *LmWT* infection compared to *LmCen*^*-/-*^ infection. Indole-3-acetaldehyde is catabolized from tryptophan through an interaction in which the Interleukin-4 induced gene 1 (IL4I1) is the rate-limiting enzyme (Rannug, 2022; Sadik et al., 2020; Zhang et al., 2020c). Indole-3-acetaldehyde can then be rearranged to form 6-formylindolo[3,2-*b*]carbazole (FICZ), another endogenous ligand of AhR (Rannug, 2022; Sadik et al., 2020; Zhang et al., 2020c). The FICZ-AhR interaction through the transcription factor c-Maf leads to the expression of anti-inflammatory IL-10 cytokine, and increased expression of IL4I1 through the transcription factor RORγt, creating an environment that supports pathogenic growth (Gargaro et al., 2016; Rannug, 2022). It is likely that the elevated levels of indole-3-acetaldehyde observed in *LmWT* infection may be responsible for the production of IL-10 through the interaction of its metabolite FICZ and AhR-c-Maf. The potential role of FICZ in the production of IL-10 in *Leishmania* infection remains to be experimentally demonstrated.

The serotonin pathway of tryptophan catabolism was also found to be differentially regulated in our dataset, with melatonin only being detected in the *LmCen*^*-/-*^ immunization but not in *LmWT* infection. Melatonin is one of the main end-products of the serotonin pathway, where the rate-limiting enzyme tryptophan hydroxylase (TPH) can catalyze the conversion of tryptophan to 5-hydrxytryptophan (5-HTP) that later yields N-acetylserotonin, and serotonin (Calvo et al., 2013; Carrillo-Vico et al., 2013; Hsu and Tain, 2020). Upon its release, melatonin has been shown to induce a strong-pro-inflammatory response in the host by stimulating significant IL-12 release in dendritic cells and macrophages (Calvo et al., 2013; Cutolo et al., 1999; Lissoni, 1999). The immune-boosting properties of melatonin have led to its usage as a vaccine adjuvant (Wang et al., 2021; Zhang et al., 2020a). Thus, the elevated levels of melatonin observed in *LmCen*^-/-^ immunization may have an endogenous adjuvant activity potentiating a strong its immune response though this remains to be tested experimentally.

Collectively, our metabolomic data indicates that the *LmCen*^*-/-*^ immunization alters all three pathways of tryptophan catabolism towards inducing host-protective immune response. The binding of Kynurenine and FICZ to AhR in *LmCen*^-/-^ appear to be limited due to their low abundance, while the tryptophan is mainly utilized for the production of melatonin that may enhance a pro-inflammatory response. The roles of the FICZ-AhR interaction and melatonin in the *LmCen*^*-/-*^ are the focus of future studies.

While the metabolic analysis of *LmCen*^-/-^ revealed tryptophan metabolism affecting the immune-protection, a similar metabolic analysis in the *Centrin*-deleted *L. mexicana* (*LmexCen*^*-/-*^) strain revealed the metabolic pathways underlying immune protection are different between the two strains (see accompanying paper). *L. mexicana* is a new world *Leishmania* species that causes cutaneous leishmaniasis. We found that in *LmexCen*^*-/-*^ the Pentose Phosphate pathway being the most highly enriched pathway of the *LmexCen*^*-/-*^ vaccine immunization that enhanced the production of nitric oxide, IL-12 and IL1-β. Thus, the immunological characteristics observed in the two mutant strains may be associated with the metabolic shifts observed in the two infections. Our findings show the importance of understanding the role of metabolomics in the generation of immune responses by vaccines.

## LIMITATIONS

The results in Table 1 and Table 2, show one of the complications of untargeted metabolomics, which is the presence of multiple isoforms of the same compound that can sometimes have opposite fold change directions and create potential ambiguity in the interpretation of the results. Additionally, the annotation of the metabolites is tentative as they are based exclusively on the formula with no additional information regarding the compound structures.

## METHODS

### Mouse strains and parasites

The animal protocol for this study has been approved by the Institutional Animal Care and Use Committee at the Center for Biologics Evaluation and Research, US Food and Drug Administration (FDA) (ASP 1995#26). In addition, the animal protocol is in full accordance with “The guide for the care and use of animals as described in the US Public Health Service policy on Humane Care and Use of Laboratory Animals 2015.” Female 6- to 8-week-old C57Bl/6 (Jackson labs) mice were immunized or infected, in the ear dermis with 1×10^6^ total stationary phase promastigotes of *LmCen*^-/-^ or *LmWT* parasites, by intradermal needle injection, in 10μl PBS.

*L. major* Friedlin (FV9) parasites were routinely passaged by inoculation into the footpads of BALB/c mice. *L. major* parasites isolated from infected lesions were cultured at 27°C in M199 medium (pH 7.4) supplemented with 10% heat-inactivated fetal bovine serum, 40 mM HEPES ((4-(2-hydroxyethyl)-1-piperazineethanesulfonic acid) (pH 7.4), 0.1mM adenine, 5mg/L hemin, 1mg/L biotin, 1mg/L biopterin, 50U/ml penicillin, and 50μg/ml streptomycin.

### *In vivo* infection

C57Bl/6 mice (n≥4 per group) that were age matched were inoculated intradermally in the ear dermis with 1×10^6^ *L. major* or *LmCen*^-/-^ promastigotes in the stationary phase. After 7 days, the ipsilateral and contralateral infected ears and the naïve ears were collected and processed for mass spectrometry.

### Mass spectrometry

The ear tissue was collected, snap frozen, and processed for mass spectrometry analysis according to SOP 7 of the Laboratory Guide for Metabolomics Experiments (Manchester). Samples were then incubated with 500 ul of 100% MeOH and sonicated. The tissue was weighed and homogenized at 40 mg/mL of 50% MeOH solution for 3 cycles in a homogenizer with Precellys. The supernatant was collected, dried down, and reconstituted in ½ of the original volume in 5 % MeOH with 0.1 % formic acid.

Untargeted analysis was performed on a Thermo Orbitrap LTQ XL with HPLC separation on a Poroshell 120 SB-C18 (2 × 100 mm, 2.7 μm particle size) with an WPS 3000 LC system. The gradient consisted of solvent A, H2O with 0.1 % Formic acid, and solvent B 100 % acetonitrile at a 200 μL/min flow rate with an initial 2 % solvent B with a linear ramp to 95 % B at 15 min, holding at 95% B for 1 minutes, and back to 2 % B from 16 min and equilibration of 2 % B until min 32. For each sample, 5 μL were injected and the top 5 ions were selected for data dependent analysis with a 15 second exclusion window.

For feature selection in the untargeted results analysis, including database comparison and statistical processing, samples were analyzed in Progenesis QI, and the pooled sample runs were selected for feature alignment. Anova p-value scores between the groups were calculated with a cutoff of < 0.05. With database matching using the [Human Metabolome Database], selecting for adducts M+H, M+Na, M+K, and M+2H and less than 10 ppm mass error, unique features were tentatively identified as potential metabolites.

### Statistical analysis of mass spectrometry datasets

Peak intensity data tables obtained from the mass spectrometry analysis were formatted into comma-separated values (CSV) files to conform to MetaboAnalyst’s requirements. The data tables were uploaded onto the one-factor statistical analysis module, and all CSV files passed MetaboAnalyst’s internal data integrity check. The data filtering was performed based on interquartile range, while sample normalization was performed based on the median of the data and the auto-scaling option was chosen to perform data scaling. However, no transformation of the data was performed. Partial least-squares discriminant analysis (PLS-DA) was performed as the dimensionality reduction method, while the Cross-validated sum of squares (Q2) performance measures were used to determine if PLS-DA models were overfitted. The significant, differentially regulated metabolites were visualized by volcano plots with cutoffs of < 0.05 false-discovery rate (FDR) and > 2-fold change (FC).

### Pathway analysis of mass spectrometry datasets

We have used two different techniques in order to identify enriched pathways in our data sets. First, we used the Functional Analysis Module (MS peaks to pathways) in MetaboAnalyst 5.0 (Pang et al., 2022; Xia et al., 2009). Detected peaks (mass-to-charge ratios + retention times) from positive and negative analytical modes of the mass spectrometer for each sample were organized into four column lists along with calculated p-values and t-scores from univariate t-tests. These peak list profiles were uploaded to the functional analysis module and passed the internal data integrity checks. The ion mode in MetaboAnalyst was set to the appropriate type depending on the analytical mode that was used to generate the data. For each analysis the mass tolerance was set to 10 ppm, the retention time units were set to minutes, and the option to enforce primary ions was checked. In parameter settings, the mummichog algorithm (version 2.0) and the modified gene set enrichment algorithm were used for all analyses. The p-value cutoff for the mummichog algorithm was left at the default (top 10% of peaks). Currency metabolites and adducts were left at default settings. Lastly, the Kyoto Encyclopedia of Genes and Genomes (KEGG) pathway library for *Mus musculus* was selected as the metabolic network that the functional analysis module would use to infer pathway activity and predict metabolite identity; only pathways/metabolite sets with at least three entries were allowed.

To confirm the enriched pathways identified with MetaboAnalyst, we also used the Ingenuity Pathway Analysis (IPA) software. Metabolite matches to the detected peaks from database searches (HMDB and LIPID MAPS), along with calculated p-values and fold changes were uploaded into IPA software for core analysis. The reference set was selected as the Ingenuity Knowledge Base (endogenous chemicals only) and direct and indirect relationships were considered during analysis. Settings in the networks, node type, data sources, confidence, and mutations tabs were left at default values. The species tab settings were set to mouse and uncategorized (selecting uncategorized species for metabolomics is necessary in IPA as most metabolites are not unique to any one species). Lastly, in the tissues and cell lines tab all tissues were considered in the analysis, whereas all cell lines were excluded from consideration. The p-value cutoff for every analysis was set as 0.05 and the fold change cutoffs were adjusted to obtain between 200-1000 analysis-ready molecules and then kept the same across different analyses.

### Metscape Analysis

The Metscape 3.1.3 App (Karnovsky et al., 2012) from the Cytoscape 3.9.1 software was used in order to build integrative network analysis of our in-vivo dataset using Metscape’s internal database which incorporates KEGG and EHMN data. The IDs of the metabolomic dataset were converted from HMDB IDs to KEGG IDs recognized by the Cytoscape software via the Chemical Translation Service (CTS) and verified by the Metaboanalyst Compound ID Conversion tool. The metabolomic dataset was then uploaded as a compound file to Metscape and the P. Value and FC Ratio cutoff points were set at 0.05 and 1.0 respectively. A Compound-gene network was created for Tryptophan Metabolism detected in our dataset, allowing us to visualize the integrated relationship among the metabolites and genes involved in this pathway.

### *In vitro* cell culture and infection

Bone marrow-derived Dendritic Cells (BMDCs) were obtained from the femurs of C57Bl/6 mice. After isolation, the bone marrow cell suspension was cultured in 6-well plates, at a concentration of 2.0×10^6^ cells per well, with complete RPMI medium supplemented with IL-4 and GM-CSF at 37°C with 5% CO_2_ for 7 days until differentiation was complete. The differentiated BMDCs were then infected overnight with stationary phase *LmCen*^*-/-*^ promastigotes at a 1:10 ratio of DCs to parasites, while the naïve controls were treated with complete RPMI media alone. The next day, the extracellular parasites were removed by washing with PBS and new media was applied. Some of the groups were treated with 600μM concentration of 1-methyl tryptophan (L-isomer, Sigma-Aldrich) or 50μM concentration of kynurenine (L-isomer, Sigma-Aldrich). After a 24hrs incubation, the cells were scraped from the wells for RNA extraction and RT-PCR.

## Graphical abstract

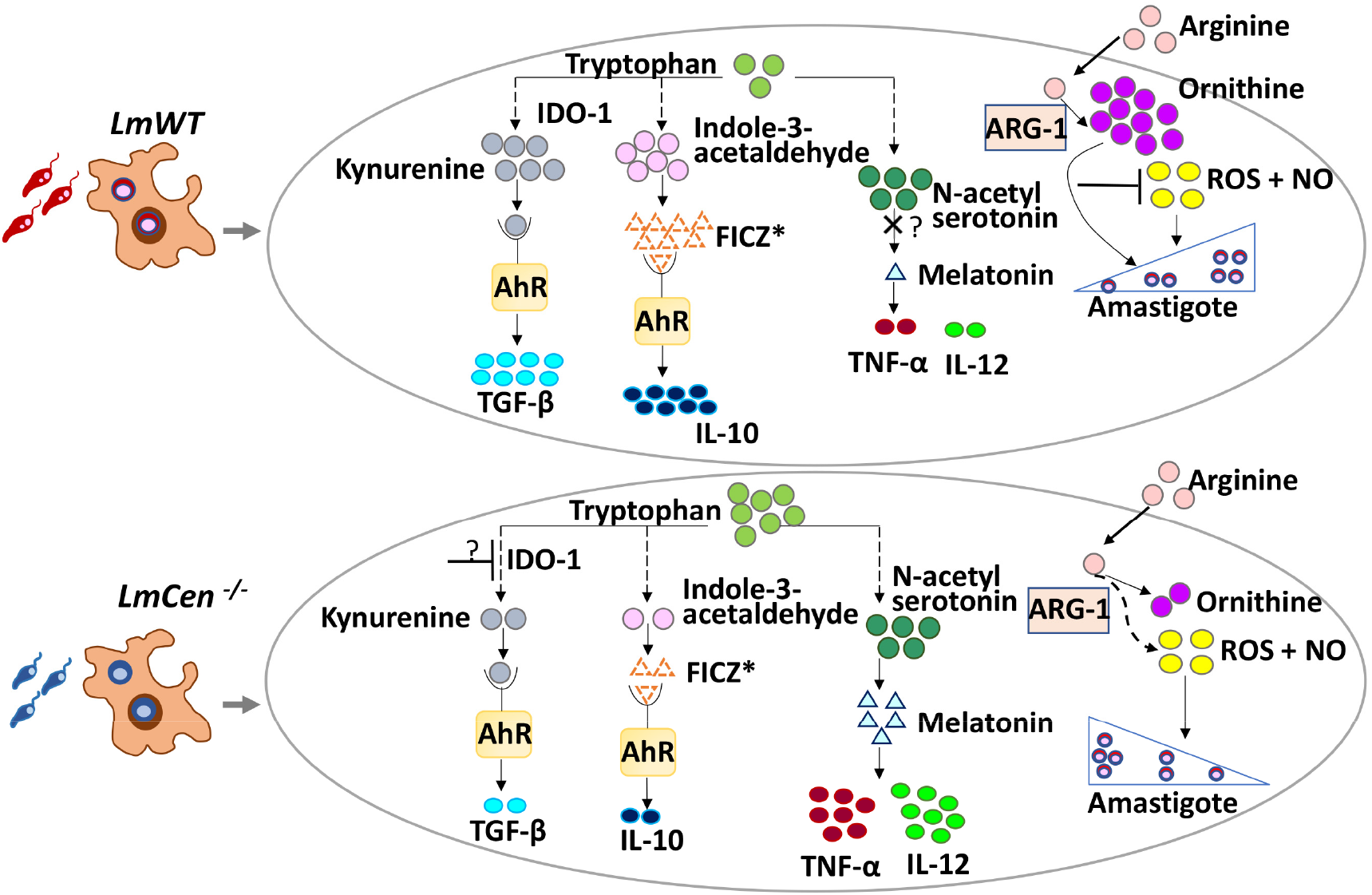

**Supp Fig 1.**
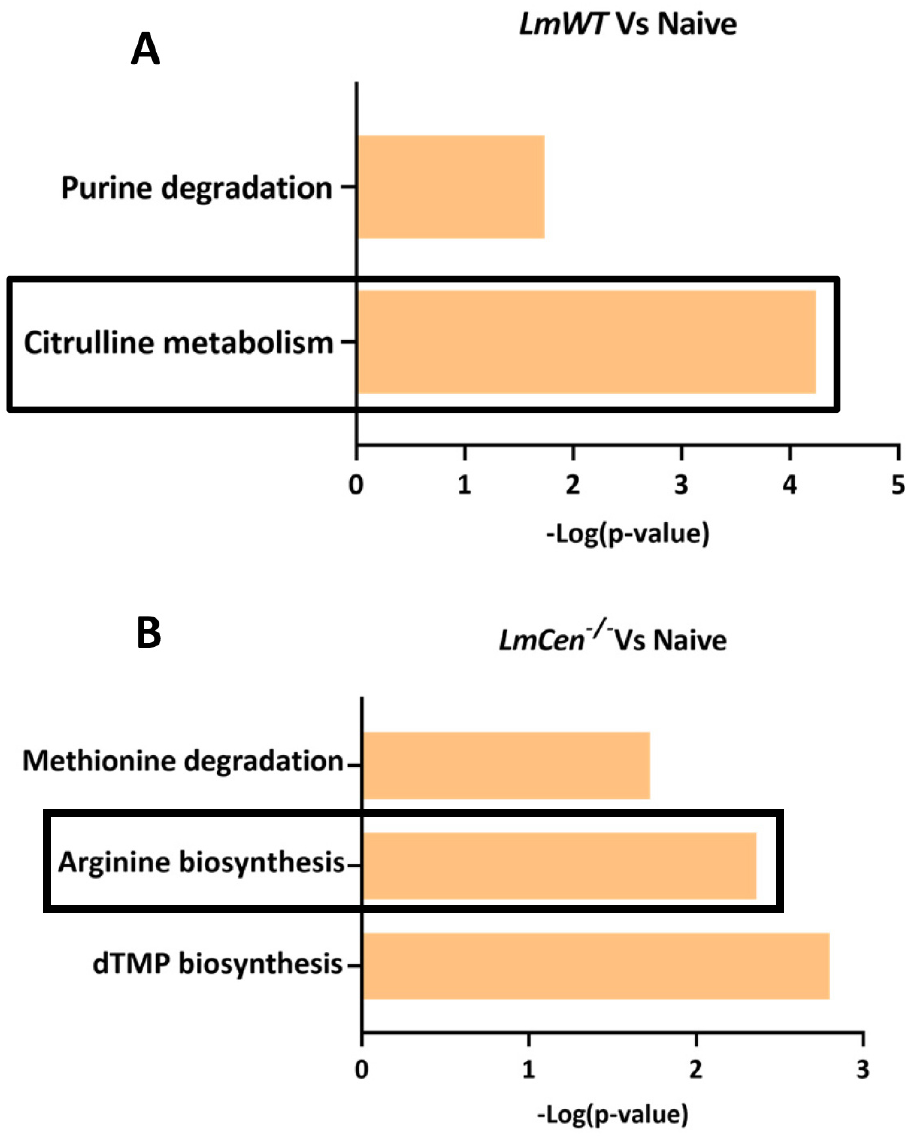

## Notes

### Competing Interest Statement

The authors have declared no competing interest.

